# Cortical astrocyte activation triggers meningeal nociception and migraine-like pain

**DOI:** 10.1101/2025.02.08.637109

**Authors:** Dara Bree, Jun Zhao, Jennifer Stratton, Dan Levy

## Abstract

Migraine attacks are believed to originate in the brain, but the exact mechanisms by which the brain generates peripheral nociceptive signals that drive migraine pain remain unclear. Sensory cortex hyperexcitability has been observed consistently across different migraine subtypes. Astrocytes detect aberrant increases in cortical activity via their Gq-coupled receptors and respond by releasing gliotransmitters and other factors with proinflammatory and nociceptive properties. In the present study, we used a rat model to investigate whether heightened cortical astrocyte Gq-coupled signaling is sufficient to drive peripheral trigeminal meningeal nociceptive responses linked to the generation of migraine headaches. We used an AAV-based chemogenetic approach that allows selective activation of cortical astrocyte Gq-GPCR signaling. We targeted astrocytes in the visual cortex as hyperexcitability in this region has been implicated in migraine. Furthermore, the meninges overlying the visual cortex are densely innervated by nociceptive fibers. We combined this chemogenetic approach with in vivo single-unit recording of meningeal nociceptors to assess changes in their ongoing activity and mechanosensitivity, along with testing of migraine-like behaviors. We further targeted calcitonin gene-related peptide (CGRP), using a monoclonal antibody (anti-CGRP mAb), to assess the relevance of cortical astrocyte activation to migraine. We discovered that heightened activation of Gq-coupled signaling in visual cortex astrocytes drives persistent discharge and increased mechanosensitivity of meningeal nociceptors. Cortical astrocytic activation also generated cephalic mechanical pain hypersensitivity, reduced exploratory behavior, and anxiety-like behaviors linked to migraine headaches. Blocking calcitonin gene-related peptide signaling suppressed astrocyte-mediated increases in meningeal nociceptor discharge and alleviated associated migraine-related behaviors. Our findings reveal a previously unappreciated role for augmented visual cortex astrocyte signaling as a triggering factor sufficient to generate meningeal nociception and migraine pain and greatly expand our understanding of migraine pathophysiology.

## Introduction

Migraine is one of the most common neurological diseases and a leading cause of global disability ^1^. A large body of work suggests that migraine headache genesis involves the activation of the trigeminal nociceptive sensory pathway that innervates the cerebral meninges ^2^. However, the early events that trigger this pain pathway during a migraine attack are poorly understood. A key migraine hypothesis implicates sensory, particularly visual, cortex hyperexcitability and related cortex-to-meninges signaling in the generation of meningeal nociception and migraine headaches ^3^. However, this hypothesis is primarily based on studies of cortical spreading depolarization (CSD), a pathophysiological event characterized by excessive cortical neuron discharge that occurs only in a subset of migraine attacks accompanied by visual and other sensory auras ^4^. Visual cortex hyperexcitability has also been consistently observed in migraine without aura ^4–7^. However, it remains unclear whether cellular events associated with increased visual cortex excitability, independent of CSD, are sufficient to generate meningeal nociception and drive migraine headaches.

Astrocytes sense and respond to neurotransmitter release during increased cortical activity ^8,9^. Neurotransmitter spillover associated with cortical hyperexcitability in migraine ^7^ can promote heightened and prolonged activation of astrocyte G protein-coupled receptors ^10–12^. Excessive activation of astrocytic Gq-coupled signaling promotes the upregulation and release of numerous astrocytic factors with algesic properties (e.g., prostanoids, cytokines, chemokines, ATP) ^8,13^. The efflux or glymphatic/lymphatic transport of these mediators into the cranial meninges may directly or indirectly activate meningeal sensory fibers and promote meningeal nociception and migraine pain.

To test whether increased astrocytic activity in the visual cortex, linked to neuronal hyperexcitability, is sufficient to generate meningeal nociception and migraine-like behaviors, we employed an adeno-associated viral (AAV)-based chemogenetic approach to selectively activate cortex astrocytes Gq-coupled signaling ^14,15^. We combined this chemogenetic tool with in vivo single-unit recording of trigeminal meningeal nociceptors to assess changes in their ongoing activity and mechanosensitivity. We further tested the development of migraine-like behaviors to determine relevance to migraine pain. Finally, we targeted CGRP signaling, implicated in migraine pathophysiology, to assess its contribution to the pro-nociceptive effects of cortical astrocyte activation. We report that stimulation of visual cortex astrocyte Gq-coupled signaling, which does not produce CSD, is sufficient to drive CGRP-mediated meningeal nociception and migraine-related pain behaviors.

## Results

### Chemogenetic activation of visual cortex astrocytes increases the firing rate of meningeal nociceptors

To determine whether increased activation of cortical astrocyte Gq-coupled signaling is sufficient to drive meningeal nociception, we selectively stimulated this astrocytic pathway using a chemogenetic approach involving the Gq-coupled Designer receptor hM3Dq ^16^ while recording single-unit activity of trigeminal nociceptors with nerve endings localized to the meninges above the primary visual cortex. Cortical astrocytes were transduced to express hM3Dq via a stereotactic injection of an AAV encoding the hM3Dq fused to mCherry under the control of the astrocyte GFAP promoter (AAV2/5-GFAP-hM3Dq-mCherry). We confirmed, using immunohistochemistry (Figure 1A-C) hM3Dq expression with high specificity to astrocytes (95.5 ± 0.9% of the hM3Dq-mCherry expressing cells were GFAP-positive) and penetrance (86.8 ± 2.2% of the GFAP-positive cells expressing hM3Dq-mCherry) in agreement with previous studies ^14,15,17–19^. We then tested astrocyte-driven nociceptor discharge in 29 meningeal nociceptors in both sexes. CNO administration triggered increased nociceptor discharge similarly in males (4/7 Aδ, 5/11C) and females (3/4 Aδ, 4/7 C), Figure 1F). Increased nociceptor activity developed in both sexes after a similar delay [males, 20 (10-27.5) min; females, 15 (10-25) min, Figure 1G]. The duration of the increased nociceptor ongoing activity was also sex-independent [males, 45 (25-47.5) min, females 45 (40-65) min, Figure 1H]. The magnitude of the increased nociceptor activity was also sex-independent [males, 1.8 (1.3-2.4) fold vs. 2.03 (1.69-2.42) fold in females, Figure 1I]. CNO administration in control hM3Dq^(-)^ rats did not affect meningeal nociceptor discharge rate [0/9 afferents, 5Aδ, 4C nociceptors from 4 males and 5 females, Figure 1F)].

**Figure 1:**
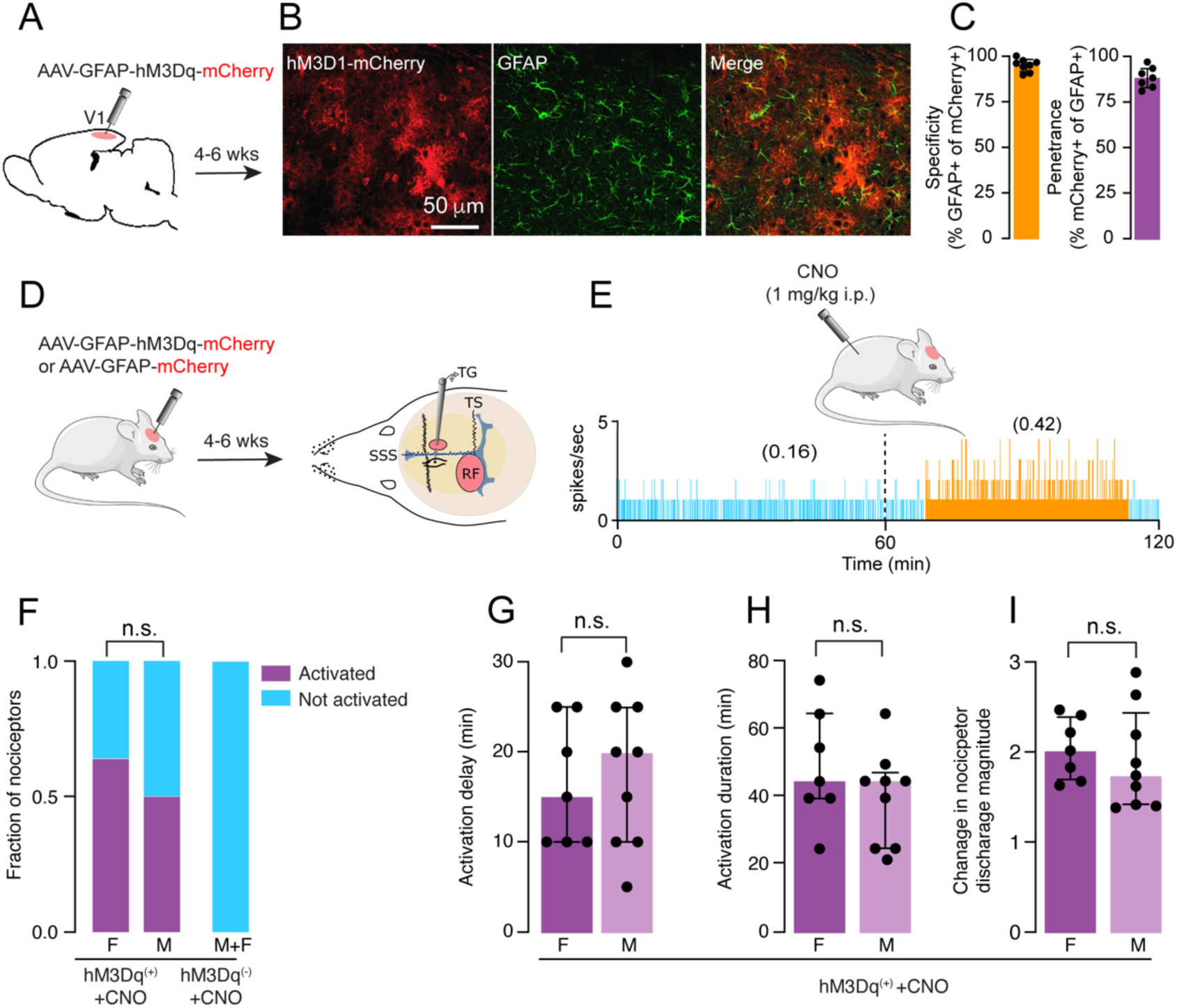
Chemogenetic stimulation of visual cortex astrocyte Gq-couple signaling activates meningeal nociceptors: **(A-C)** Cortical injection of AAV5-GFAP::hM3Dq-mCherry led after 4 weeks to hM3Dq-mCherry expression (red) in astrocytes (green) with high very high specify (>94% of mCherry positive cells were also GFAP positive, n=3) and high penetrance (>88% of GFAP positive cells were also mCherry positive, n=3). Scale bar is 50mm. **(D)** Experimental design for single-unit recordings of meningeal nociceptor responses following cortical astrocyte stimulation. For electrophysiological recordings, two skull openings (red ovals) were made. Meningeal nociceptor activity was recorded from the left trigeminal ganglion (TG) using a tungsten microelectrode inserted through a craniotomy over the contralateral hemisphere. An ipsilateral craniotomy was made to expose a part of the left transverse sinus (TS) and its vicinity to find the nociceptor’s receptive field (RF). **(E)** An example of a two-hour experimental trial depicting the activity of an Aδ meningeal nociceptor at baseline (light blue) and the increased discharge (orange) observed following injection of CNO (1 mg/kg, i.p.) at 1 h in hM3Dq^(+)^ rats to activate cortical astrocytes chemogenetically. Afferent discharge rates in spikes/sec are shown in parentheses. **(F)** The propensity of meningeal nociceptors to become activated following CNO injection in hM3Dq^(+)^ rats and control hM3Dq^(-)^ animals. Data is presented for females and males in the experimental group and the combined data from control males and females. Meningeal nociceptors recorded in females and males following astrocyte activation had similar activation propensities (p > 0.05, Chi-square test). **(G-I)** The magnitudes of the increased nociceptor activity, the latency of their development, and their durations were similar in males and females. Data is shown for individual nociceptors recorded in female and male rats with the median and interquartile range (p > 0.05, Mann-Whitney test).

To exclude the possibility that the astrocyte-driven meningeal nociceptor response resulted from the triggering of a CSD event, we recorded changes in DC potential and CBF in hM3Dq^(+)^ rats in response to CNO administration. We observed no astrocyte-driven changes in vascular or neural changes suggestive of CSD elicitation (Females, 0/4 experiments; males, 0/4 experiments, Supplementary Figure S1), which is in agreement with previous studies ^14,15,20,21^.

### Cortical astrocyte activation drives mechanical sensitization of meningeal nociceptors

The development of augmented mechanical responsiveness (i.e., mechanical sensitization) of meningeal nociceptors is thought to underlie the exacerbation of migraine headaches during normally innocuous physical activities, such as coughing and physical movement, which cause momentary increases in intracranial pressure and meningeal deformation ^2,22,23^. We, therefore, asked next whether activation of cortical astrocytes also enhances the mechanosensitivity of meningeal nociceptors. Following CNO administration, sensitization developed in both sexes with a similar propensity [Females, 7/11 (3/4 Aδ and 4/7 C); males, 11/18 (5/7 Aδ and 6/11 C), Figure 2C]. CNO administration in hM3Dq^(-)^ rats did not sensitize meningeal nociceptors [1/9 (1/5 Aδ; 0/4 C) from 4 males and 5 females, Figure 2C]. The latency to develop mechanical sensitization was sex-independent [females, 30 (15,45) min; males, 30 (15,30) min, Figure 2D], as was the duration of the sensitization [females, 75 (60,90) min; males, [90 (60,90) min, Figure 2E]. Meningeal nociceptors recorded in females and males also exhibited similar sensitization magnitudes [females, 2.1 (1.97-2.4) fold; males, 2.04 (1.91-2.6), Figure 2F].

**Figure 2:**
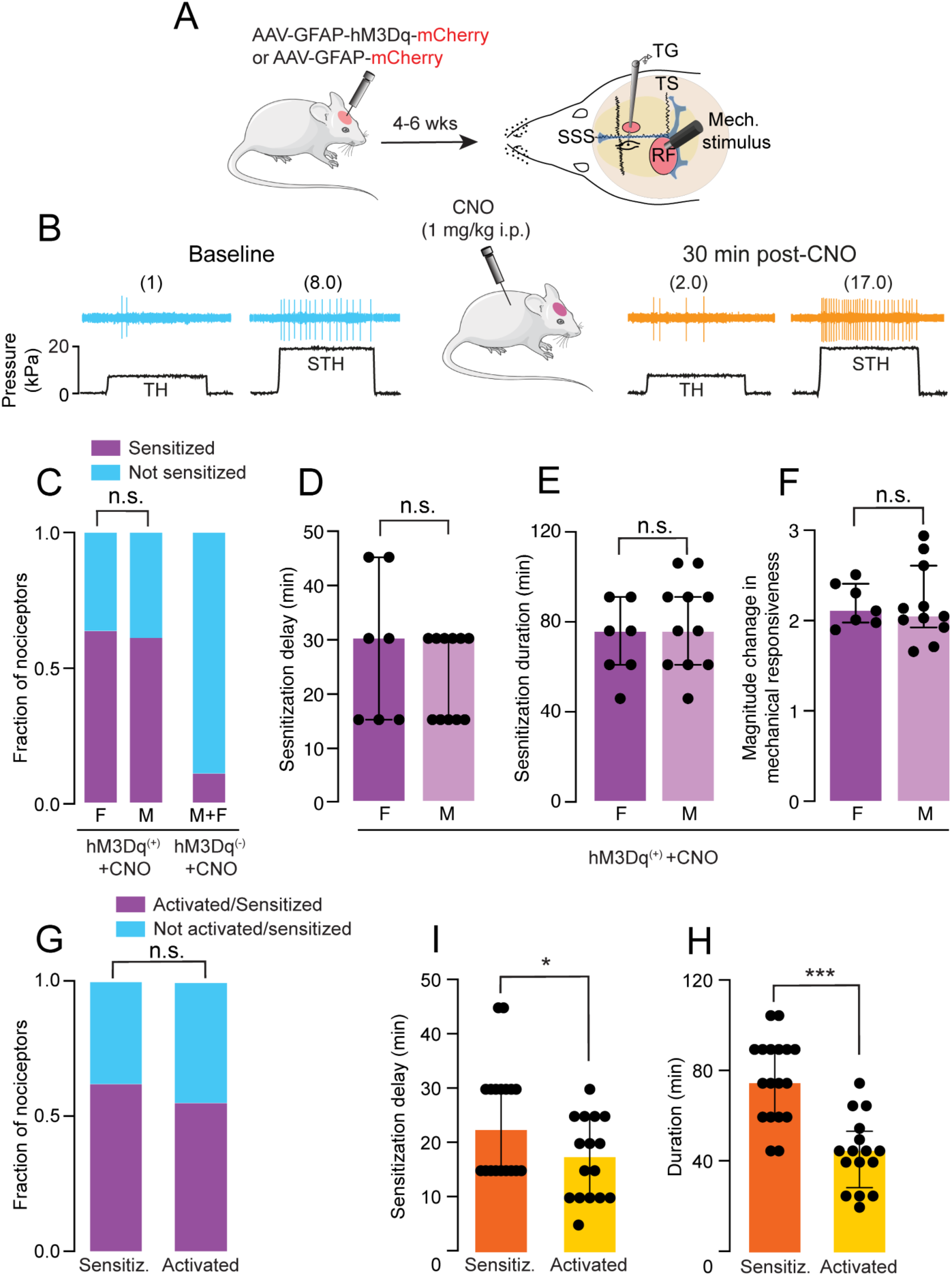
Chemogenetic activation of cortical astrocytes Gq-coupled signaling induces mechanical sensitization of meningeal nociceptors. **(A)** Experimental design for electrophysiological recording of changes in meningeal nociceptor mechanosensitivity following cortical astrocyte Gq-GPCR pathway activation. Quantitative mechanical stimuli were delivered to the nociceptors’ receptive field (RF) using a feedback-controlled mechanical stimulator**. (B)** Examples of experimental trials depicting the responses of a meningeal nociceptor (C-unit) to threshold (TH) and suprathreshold (STH) mechanical stimuli (black traces) during baseline recording (light blue traces) and then at 30 min following the administration of CNO (orange traces) to an animal injected with AAV5-GFAP::hM3Dq-mCherry 5 weeks earlier [(hM3Dq^(+)^]. Mechanically-evoked responses in spikes/sec are in parentheses. Note the increased TH and STH responses post-CNO injection. **(C)** The propensity of meningeal nociceptors to develop mechanical sensitization following CNO injection in hM3Dq^(+)^ and control animals [hM3Dq^(-)^]. Data is presented for females and males in the experimental group, and the combined data is from males and females in the control group. Meningeal nociceptors recorded in females and males following astrocyte Gq-GPCR pathway activation had similar sensitization propensities (p > 0.05, Chi-square test). **(D-F)** The magnitudes of the increased nociceptor mechanosensitivity, the latency of their development, and durations were similar in males and females. Data is presented for individual nociceptors with the median and interquartile range (p > 0.05, Mann-Whitney test). (**G)** The nociceptors’ activation and sensitization propensities following CNO injection in hM3Dq^(+)^ rats were not different (p > 0.05, Chi-square test). **(H-I)** Mechanical sensitization developed with a longer latency than persistent activation and had a longer duration when compared to the duration of nociceptor activation. Data shows individual nociceptor responses and the median with interquartile range (latency differences, p < 0.0001 Mann-Whitney test, duration differences, p < 0. 5, Mann-Whitney test).

In previous studies, mechanical sensitization of meningeal nociceptors was independent of the prolonged increase in their ongoing discharge ^24,25^. Here, we found a similar incidence of persistent activation and mechanical sensitization in response to chemogenetic stimulation of cortical astrocytes (18/29 vs. 16/29, Figure 2G). However, mechanical sensitization developed with a longer latency than the persistent activation [22.5 (15,30) min vs. 17.5 (10,25) min, Figure. 2I]. The duration of the mechanical sensitization was also longer when compared to the duration of the nociceptor activation [75 (60,90) min vs. 45 (28.75,53.75) min), Figure 2H], pointing to different mechanisms underlying meningeal nociceptors’ activation and mechanical sensitization in response to cortical astrocyte activation.

### Cortical astrocyte activation drives migraine-like pain behaviors

Cephalic cutaneous mechanical hypersensitivity is a key sensory finding in migraine attributed to the activation of meningeal nociceptors and the ensuing sensitization of dorsal-horn neurons that receive convergent sensory input from the meninges and cephalic skin. We, therefore, asked next whether activating cortical astrocytes also drives cephalic mechanical pain hypersensitivity. Following astrocyte stimulation, we observed reduced mechanical pain thresholds for at least 4 h post-CNO administration in both sexes that recovered to baseline values at 24 h (Figure 3B, C). The astrocyte-related reduction in mechanical thresholds in hM3Dq^(+)^ animals was accompanied by increased pain response that was also resolved at 24 h (Figure 3D, E). CNO injection in control hM3Dq^(-)^ animals did not affect cephalic pain thresholds and related pain behaviors (Figure 3B-E).

**Figure 3:**
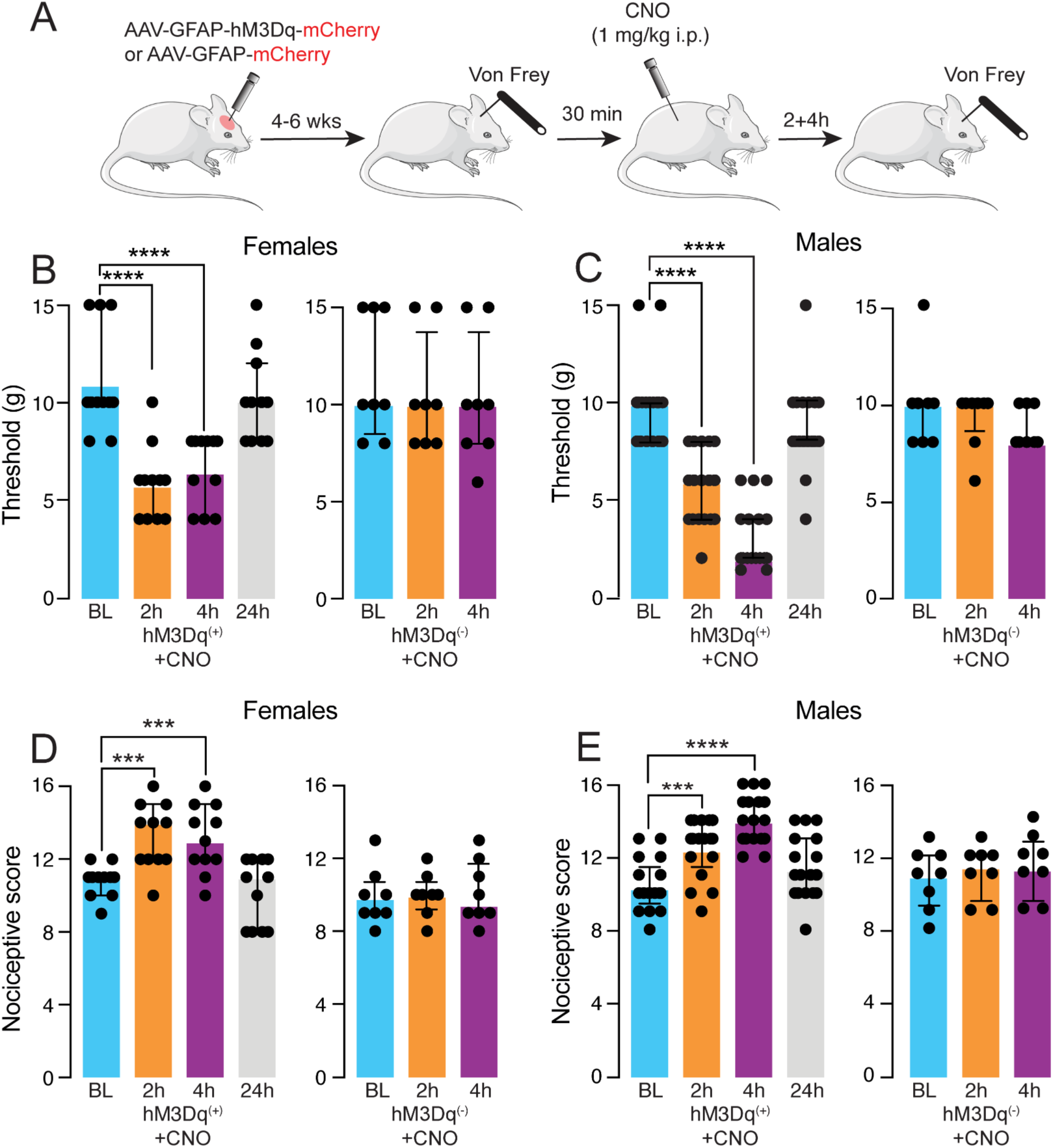
Gq pathway activation in cortical astrocytes drives cephalic cutaneous hypersensitivity. **(A)** Experimental schematics. **(B, C)** Time course changes in cephalic mechanical pain thresholds following CNO injection in females (n = 11) and males (n =17) hM3Dq^(+)^ and control hM3Dq^(-)^ rats (female, n = 8; male n = 8). Decreased thresholds were observed in both sexes at 2 and 4 h post-CNO, with recovery at 24 h (q < 0.001, Friedman test with post hoc tests). CNO injection in females and males hM3Dq^(-)^ did not change cephalic mechanical thresholds (Females and males, p > 0.5 Friedman test). **(D, E)** Time course changes in pain response to cephalic mechanical in females and males hM3Dq^(+)^ and control hM3Dq^(-)^ rats. Increased pain scores were observed in both sexes at 2 and 4 h post-CNO, with recovery at 24 h (Females and males, q < 0.001, Friedman test with post hoc tests). CNO injection in females and males hM3Dq^(-)^ rats did not affect cephalic mechanical thresholds (Females and males, p < 0.05 Friedman test with post hoc tests).

Decreased exploratory locomotion and reduced rearing in the open-field test have been linked to increased meningeal nociception related to migraine pain ^26,27^. In the open-field test, reduced exploratory behavior in the middle zone - measurement of increased anxiety - has also been linked to migraine mechanisms ^28^. We, therefore, asked next whether cortical astrocyte activation leads to deficits in exploratory behavior and anxiety-like behavior. Because open-field behaviors exhibit large inter-animal variability, we treated animals as their own controls and compared behaviors observed 4 h following CNO injection to those observed at baseline before CNO injection. Repeated open-field testing using the paradigm above can lead to behavioral changes, likely due to habituation. Therefore, we compared the behavioral changes in response to CNO treatment in hM3Dq^(+)^ rats with those observed in control hM3Dq^(-)^ rats. We observed a significant decrease in overall locomotion following astrocytic activation in females and males (Figure 4B). Activating cortical astrocytes also decreased rearing behavior in both sexes (Figure 4C). A reduction in center zone explorative behavior was observed in females, and there was a strong trend towards increased anxiety-like behavior also in males (Figure 4D).

**Figure 4:**
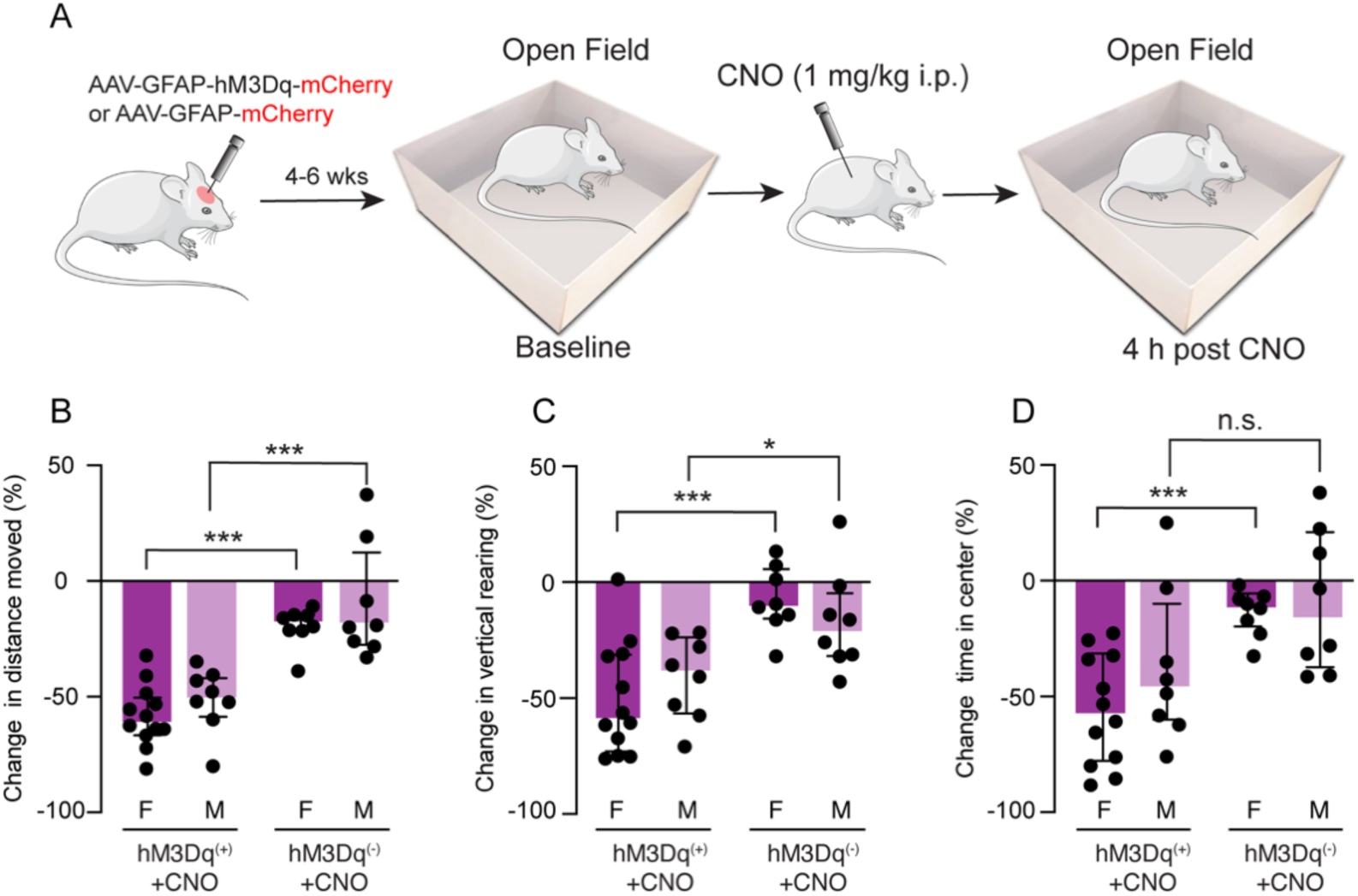
Gq pathway activation in cortical astrocytes leads to deficits in open-field activity. **(A)** Experimental schematics. **(B)** Changes in total distance moved in hM3Dq^(+)^ and control hM3Dq^(-)^ rats 4 h following CNO injection. Data is shown for individual rats and the median with interquartile range (females n = 12, males n=8; p < 0.001, Mann-Whitney test). **(C)** Change in vertical rearing in hM3Dq^(+)^ and control hM3Dq ^(-)^ rats following CNO injection. Data is shown for individual rats with the median and interquartile range (females n = 12, p < 0.001, Males n = 8, p < 0.05, Mann-Whitney test). **(D)** Changes in center time duration in hM3Dq^(+)^ and control hM3Dq^(-)^ rats following CNO injection. Decreased time indicates anxiety behavior. Data is shown for individual rats and the median with interquartile range (Females (n=12) p < 0.001, Males (n =8) p > 0.05, Mann-Whitney test).

### Anti-CGRP mAb inhibits the astrocyte-driven activation of meningeal nociceptors

Peripheral CGRP signaling is considered a key mechanism underlying migraine pain. However, the role of CGRP in mediating migraine-related activation and sensitization of meningeal nociceptors remains unclear ^2^. We, therefore, asked if blocking CGRP signaling using a clinically relevant anti-CGRP monoclonal antibody (mAb) could inhibit the astrocyte-driven meningeal nociceptor responses. Anti-CGRP mAb prevented the astrocyte-related meningeal nociceptor activation when compared to pretreatment with the isotype control mAb [2/17 nociceptors (1/10 females; 1/7 males) vs. 8/16 nociceptors (4/7 females; 4/9 males); Figure 5B)]. To test whether CGRP’s role in mediating the astrocyte-driven meningeal nociception is sex-dependent, we compared the data obtained from anti-CGRP mAb-treated animals with a combined dataset collected from the isotype control and non-treated animals (which had similar activation propensities). This analysis revealed that meningeal nociceptor activation following astrocyte activation was CGRP-dependent in both sexes [females (1/10 vs. 11/18 nociceptors; males (1/7 vs. 17/27); Figure 5C]. The anti-CGRP mAb failed to inhibit the astrocyte-related meningeal nociceptor mechanical sensitization. In rats treated with the anti-CGRP mAb sensitization developed in 9/17 nociceptors (4/10 females; 5/7 males) and 12/16 nociceptors (5/7 females; 7/9 males) following treatment with the isotype control (Figure 5D). The anti-CGRP mAb pretreatment also did not affect any of the sensitization parameters when compared to the isotype treatment (Figure 5E-G), including the latency [30 (15,37.5) min vs 22.5 (15,30) min], duration [75 (52.5,90) min vs. 60 (45,75) min] and magnitude [2.2 (1.93,2.5) fold vs. 1.75 (1.64,2.20) fold].

**Figure 5.**
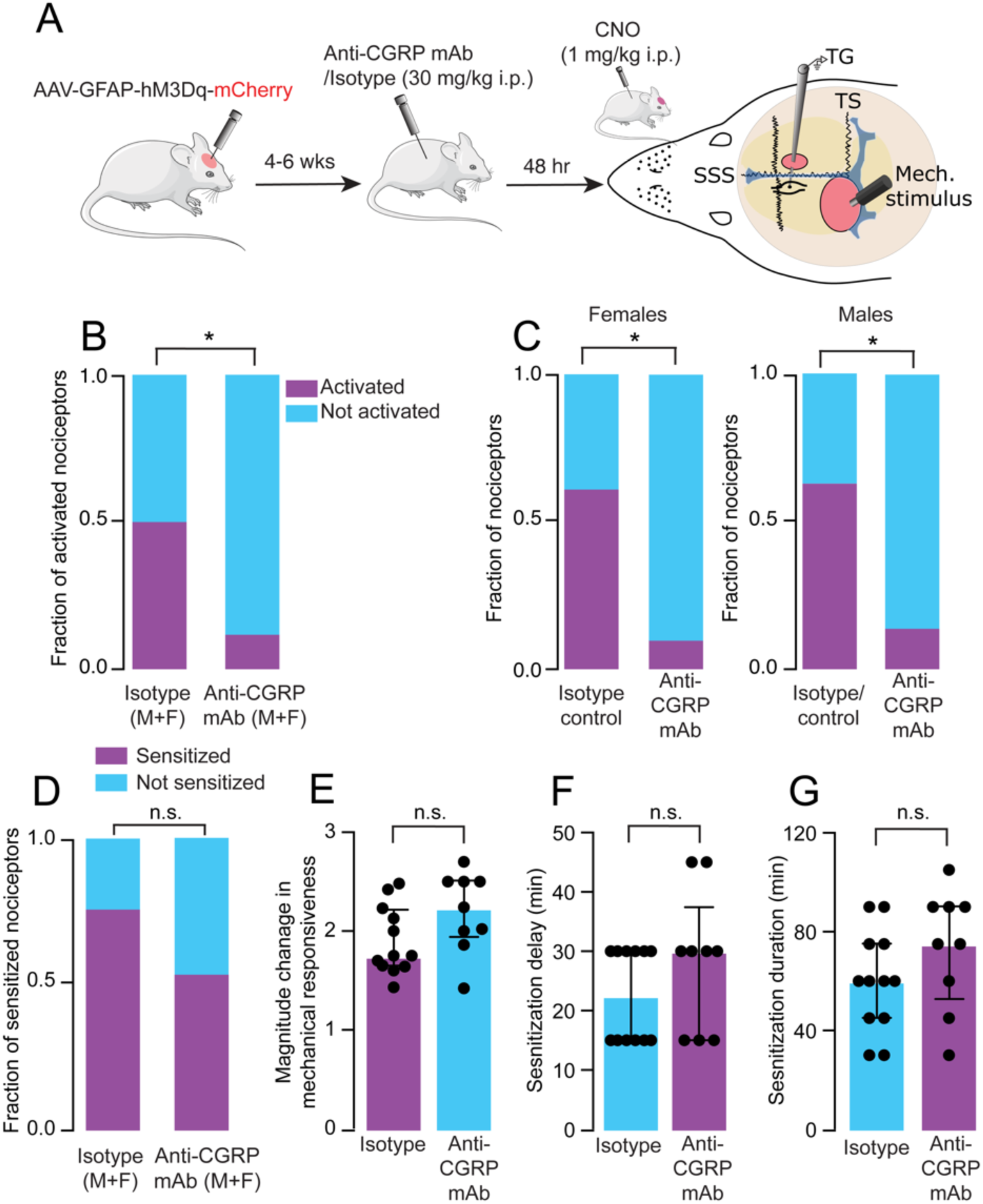
Anti-CGRP mAb antibody inhibits the activation but not mechanical sensitization of meningeal nociceptors following Gq-coupled pathway activation in cortical astrocytes. **(A)** Experimental schematics. **(B)** Targeting CGRP signaling using an anti-CGRP mAb inhibits the activation of meningeal nociceptors in response to cortical astrocyte Gq-GCPR pathway stimulation (p < 0.01, Chi-square test). **(C)** Astrocytic-evoked meningeal nociceptor activation was CGRP-dependent in females and males (p < 0.01 Chi-square test, anti-CGRP mAb vs. isotype control). **(D)** Astrocytic-driven mechanical sensitization of meningeal nociceptors was not inhibited by an anti-CGRP mAb treatment (p > 0.05, Chi-square test). **(E-G)** All sensitization parameters, including the magnitude of mechanically-evoked responses, the latency to the develop sensitization following CNO injection, and its duration, were unaffected by anti-CGRP mAb (isotype control, n = 12, anti-CGRP mAb, n = 9, p > 0.05, Mann-Whitney test).

### Anti-CGRP mAb diminishes astrocyte-mediated migraine-like cephalic mechanical hypersensitivity and deficits in open-field activity

We next asked whether anti-CGRP mAb treatment can also ameliorate the development of migraine-like cephalic pain hypersensitivity. After CNO administration, mechanical thresholds decreased at 4 h in males and females pretreated with the anti-CGRP mAb and isotype control (Figure 4B). However, the magnitude of decreased mechanical thresholds was lower in the anti-CGRP mAb treatment group compared to the isotype control group Figure 6C). Anti-CGRP mAb treatment had a similar inhibitory effect on increased pain response in both sexes (Figure 6D, E).

**Figure 6.**
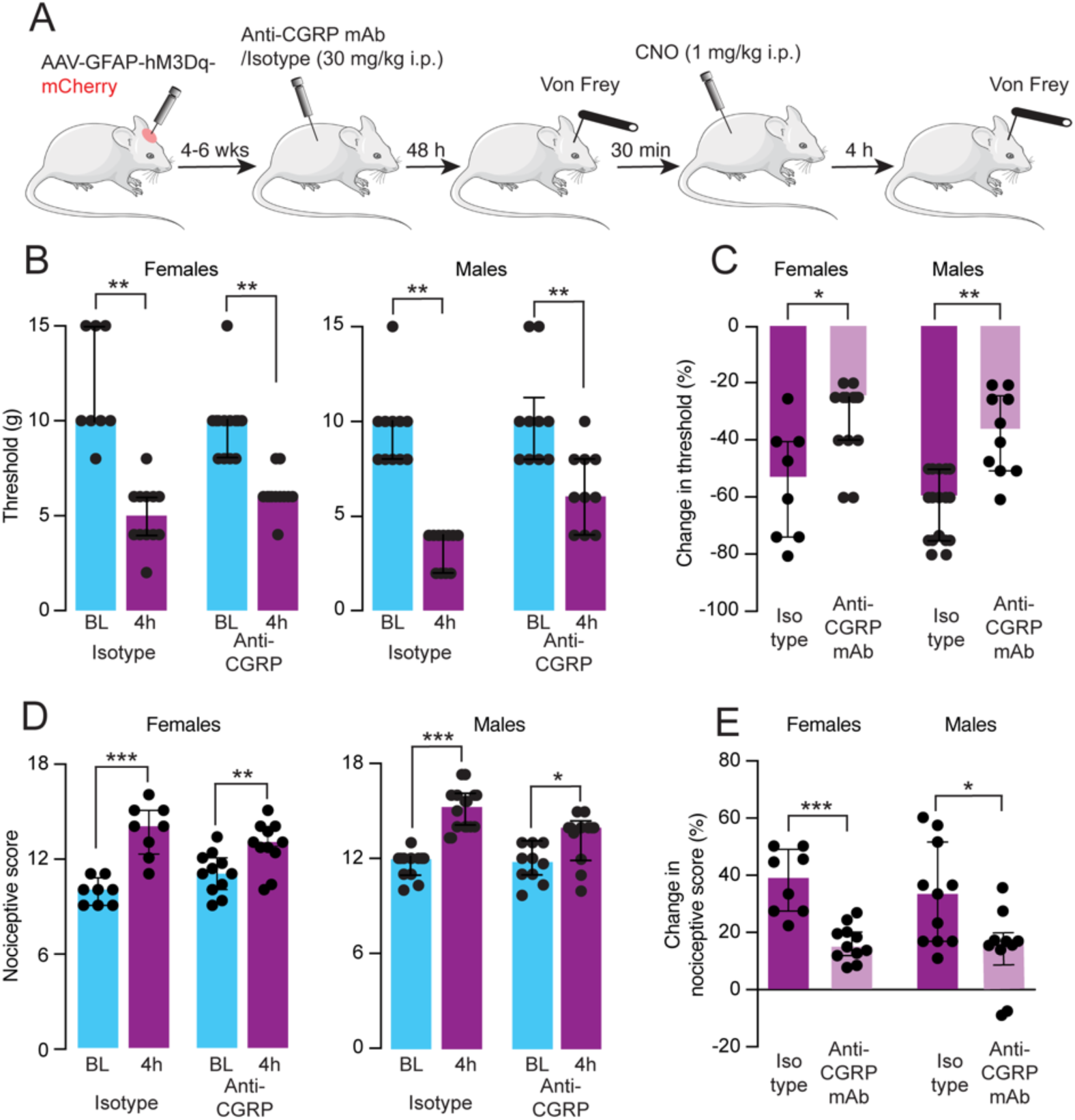
Anti-CGRP treatment inhibits cortical astrocyte-driven cephalic cutaneous pain hypersensitivity in males and females. **(A)** Experimental design. **(B)** Cephalic mechanical thresholds at baseline and 4 h following CNO injection in females and males hM3Dq^(+)^ pretreated with anti-CGRP mAb or isotype control. Data is shown for individual rats with the median and interquartile range (p < 0.01, Wilcoxon test for females, IgG n = 8, anti-CGRP n = 11 and males, IgG n = 11, anti-CGRP n = 10). **(C)** Magnitudes of change in threshold following CNO injection. Data is shown for individual rats and the median with interquartile range (females, p < 0.01; males, p < 0.05, Mann-Whitney test). **(D)** Pain scores in response to cephalic mechanical stimulation at baseline and at 4 h following CNO injection in females and males hM3Dq^(+)^ pretreated with anti-CGRP mAb or isotype control. Data is shown for individual rats and the median with interquartile range (females isotype control group, p < 0.001, Anti-CGRP mAb group, p < 0.05, males, isotype control group, p < 0.001, Anti-CGRP mAb group, p < 0.05, Wilcoxon test, n = as in B). **(E)** Magnitudes of change in pain score following CNO injection. Data is shown for individual rats and the median with interquartile range (females, p < 0.0001; males p < 0.05; Mann-Whitney test, n = as in B).

We next asked whether the astrocyte-driven deficits in open-field behaviors depended on CGRP signaling (Figure 7A). Compared to isotype control pretreatment, the anti-CGRP mAb reduced the decrease in total distance traveled in both sexes (Figure 7B). The anxiety-related reduction in center zone explorative behavior was improved only in females pretreated with the anti-CGRP mAb (Figure 7C). In contrast, the anti-CGRP mAb had no effect on the astrocyte-driven decrease in rearing in both sexes (Figure 7D).

**Figure 7.**
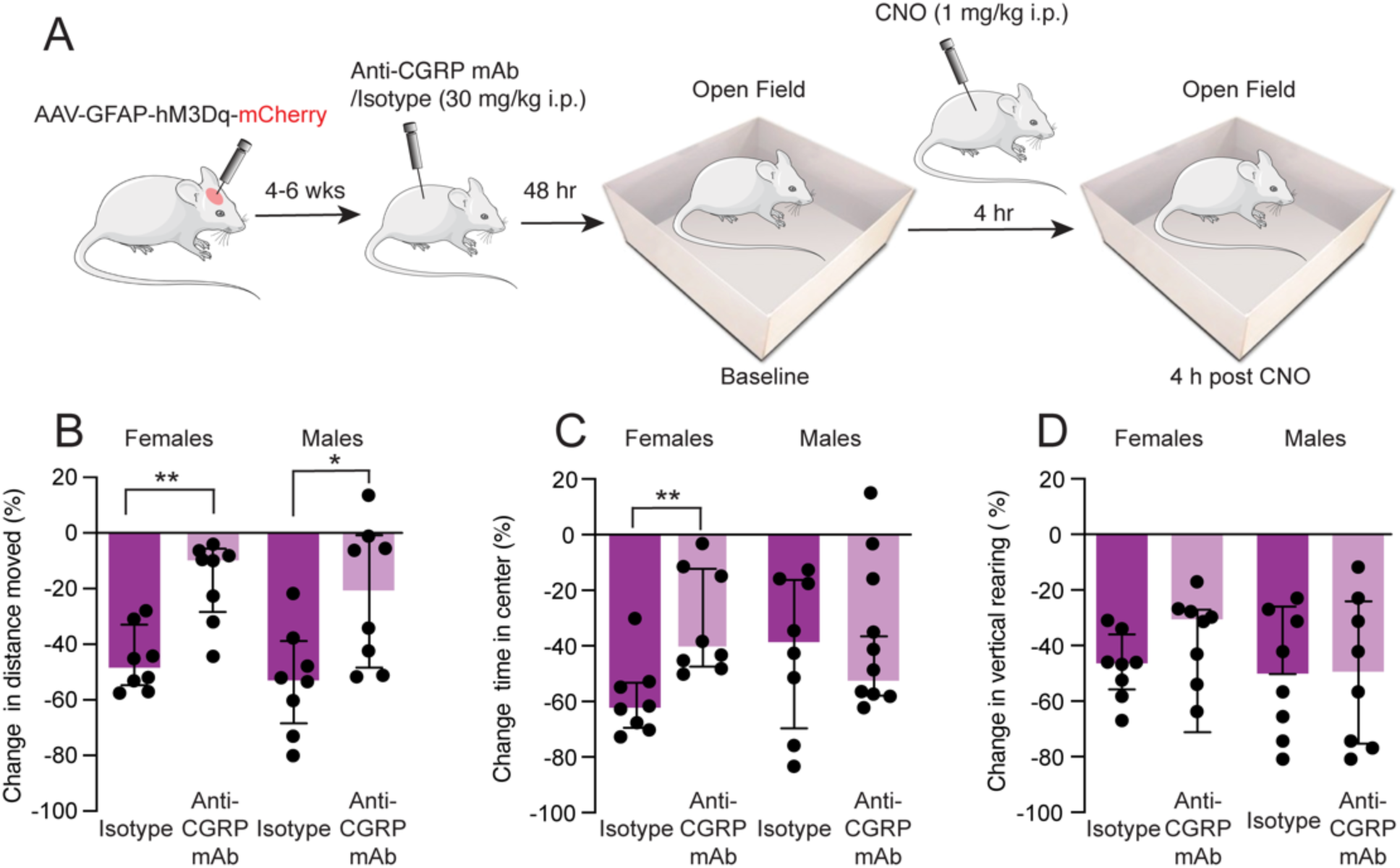
Anti-CGRP treatment inhibits cortical astrocyte-driven open-field deficits selectively in females. **(A)** Schematic of the experiments. **(B-D)** Change in open-field behavior, including total distance traveled, rearing, and time spent in the center zone, in hM3Dq^(+)^ rats injected with CNO after pretreatment with anti-CGRP mAb or isotype control. Data is shown for individual rats with the median and interquartile range. Anti-CGRP mAb pretreatment inhibited the reduction of exploratory locomotion in both sexes (p < 0.01 vs. isotype control, Mann-Whitney test, n = 8) and the time spent in the center zone only in females (p < 0.01 vs. isotype control, Mann-Whitney test, n = 8).

## Discussion

Astrocytes have been implicated in pain mechanisms. However, whether increased cortical astrocyte signaling is sufficient to generate migraine pain remains unexplored. Here, using a chemogenetic approach to manipulate cortical astrocytes, we demonstrate that activation of the Gq-GPCR signaling pathway in visual cortex astrocytes is sufficient to provoke prolonged discharge and mechanical sensitization of meningeal nociceptors, the sensory neural population whose activation is thought to mediate migraine headaches. Combining our chemogenetic tool with studies of cephalic cutaneous mechanosensitivity and open-field behaviors, we further demonstrate that activation of cortical astrocytes induces migraine-like cephalic allodynia and deficits in open-field activity reminiscent of migraine headache pain. Finally, we show that astrocyte-driven increases in meningeal nociceptor discharge and migraine-like behaviors involve CGRP signaling.

The exact astrocytic mechanism responsible for driving persistent meningeal nociceptor discharge is unclear but might be related to the prolonged Ca^2+^ elevation following the stimulation of their Gq-coupled signaling following hM3Dq activation ^15^. Prolonged Gq-coupled signaling and the ensuing Ca^2+^ elevation in cortical astrocytes could lead to the elaboration of gliotransmitters with pro-nociceptive properties, such as glutamate and ATP ^10^, that can propagate across the meninges ^29^ and stimulate meningeal nociceptors’ peripheral nerve endings. Heightened and prolonged Gq-coupled pathway activation in astrocytes can also lead to the release of inflammatory mediators such as prostaglandins ^13^ that augment the responsiveness of meningeal nociceptors ^24^. Although a local cortex-to-meninges signaling could drive the prolonged discharge and sensitization of meningeal nociceptors following the activation of cortical astrocyte Gq-coupled signaling pathway, we cannot exclude the possibility that the same cortical mediators flow via the CSF into the trigeminal ganglion and directly activate nociceptors’ somata, as suggested in the CSD model of migraine ^30^. However, the mechanisms underlying the mechanical sensitization of meningeal nociceptors seem localized to the meninges ^24,31–34^.

Cortical astrocyte stimulation produced cephalic allodynia in both sexes. A likely contributing mechanism involves the sensitization of medullary and upper cervical dorsal horn nociceptive neurons in response to the prolonged activation of meningeal nociceptors, which was also sex-independent. Notably, following cortical astrocyte activation, rats exhibited a longer duration of cephalic allodynia than in the CSD migraine model (> 4 h vs. < 2 h) ^35^, which may be related to the differences between the cortical responses associated with CSD and those occurring following the selective stimulation of visual cortex astrocytes. CSD propagates throughout the ipsilateral cortex and engages descending inhibiting pathways that can limit the development of central sensitization and dorsal horn responses evoked by stimulating cutaneous afferents ^36^. Although the activation of astrocytes can alter cortical synaptic excitatory and inhibitory balance ^14,19,37^, we targeted the visual cortex, which does not modulate cutaneous mechanical sensitivity ^36^.

Cortical astrocyte activation led to decreased open-field activity reminiscent of migraine pain in both sexes. In contrast, only females exhibited anxiety-like behavior. The development of anxiety-like behavior likely reflects sexually dimorphic processing in CNS-related anxiety circuits. The earlier finding that a single CSD event is not associated with an anxiety-like behavior ^35^ points to a mechanism involving a CNS process triggered by visual cortex astrocyte activation that is potentially unrelated to the augmented nociceptors’ responses.

Targeting CGRP signaling using mAb is an effective prophylactic treatment for migraine, but only in about half of the patients ^38^. Furthermore, blocking CGRP signaling does not always completely relieve headache and associated migraine symptoms ^39^, suggesting that not all migraine-triggering processes are CGRP-dependent. We found that anti-CGRP mAb pretreatment blocked the activation of meningeal nociceptors in both females and males but did not affect their mechanical sensitization, which may reflect the incomplete migraine prophylactic effects of anti-CGRP treatments in a subset of patients. The mechanism underlying these distinct responses to anti-CGRP mAb remains to be elucidated but may be related to the mAb therapeutic site of action.

Meningeal application of CGRP does not excite meningeal nociceptors, at least not in male rodents ^40^. However, in the CSD model, the influx of CSF-containing CGRP excites trigeminal ganglion somata in both sexes ^30^, and CGRP administration to the trigeminal ganglion produces migraine-like pain behaviors in males and females ^41^. The trigeminal ganglion may serve as the site of action for anti-CGRP mAb, which mediates the inhibition of the astrocytic-driven meningeal nociceptor activation. The sex-independent inhibitory effect of the anti-CGRP mAb on the meningeal nociceptor activation we observed is congruent with the findings that an anti-CGRP mAb inhibits meningeal nociceptor input to trigeminal dorsal horn neurons in both male and female rats ^42^. We further propose that the CGRP-independent mechanical sensitization occurs at the level of the nociceptors’ meningeal nerve endings. The finding that an anti-CGRP mAb failed to exert an anti-inflammatory effect in the meninges ^43^ is also consistent with its inability to inhibit inflammatory-driven mechanical sensitization.

Consistent with the lack of sex differences in the effect of anti-CGRP mAb on meningeal nociceptor activation, our data suggest that CGRP signaling underlies cortical astrocyte-driven cutaneous mechanical hypersensitivity and reduced open-field locomotion in both females and males. However, the astrocytic-driven decrease in rearing behavior was CGRP-independent. Because rearing behavior is likely associated with brief increases in intracranial pressure and meningeal mechanical stretching that could activate mechanosensitive nerve fibers ^23,31^, the mechanical sensitization of meningeal nociceptors, which was also CGRP-independent, could contribute to the decreased rearing. The female-specific anxiety-like behavior we observed was also CGRP-dependent, further suggesting the involvement of a sexually dimorphic CNS mechanism, which anti-CGRP mAb may also target ^44^.

In summary, we report that heightened visual cortex astrocyte Gq-coupled signaling activation is sufficient to drive meningeal nociception and migraine-like behaviors involving peripheral CGRP signaling. Further studies are needed to elucidate the molecular signaling downstream of cortical astrocyte activation, including the involvement of Ca^2+^ and molecules released from astrocytes responsible for triggering meningeal nociceptors and driving migraine pain.

## Methods

### Animals

All experimental procedures were approved by the Beth Israel Deaconess Medical Center Institutional Animal Care and Use Committee. Male and female rats (Sprague-Dawley, 220–300 g, Taconic, USA) were housed in pairs with food and water *ad libitum* under a constant 12-hour light/dark cycle at room temperature. All procedures and testing were conducted during the light phase of the cycle. Experimental animals were randomly assigned to different treatment groups. Responses to different treatments were studied in a blinded fashion.

### AAV injection

Rats were anesthetized with isoflurane and placed in a stereotaxic apparatus. An incision was made to expose the skull, and burr holes were drilled over the visual cortex (−7.5 mm AP, +/-2.6 and +/- 4.5 mm ML, with respect to bregma). a Hamilton syringe was lowered 100-300 mm below the pia, and 100 nl of AAV2/5-GFAP-hM3Dq-mCherry (7×10^12^ vg/ml) was injected at 100 nL/min. The viral prep was made using pAAV-GFAP-hM3Dq-mCherry, a gift from Bryan Roth (Addgene #50478-AAV5). In control experiments, AAV-GFAP-mCherry was microinjected (Addgene# 58909-AAV5). At the end of the injections, the skin incision was sutured, and animals were given SR meloxicam (4 mg/ml) for post-surgical analgesia.

### Immunohistochemistry

Rats were first given a lethal dose of urethane (2 g/kg) and perfused transcardially with 0.025% heparin in PBS, followed by 4% paraformaldehyde (PFA). Mice were decapitated, and heads were postfixed in 4% PFA at 4^0^ C for 24 hours. The brain was then extracted and cryoprotected with 30% sucrose (48 hours) before being embedded in OCT (Fisher Scientific), rapidly frozen over dry ice and 20 μm cryosections, cut on a cryostat (Leica) and washed 3 times with PBS. To label the hM3Dq-mCherry and astrocytes, the sections were immune-stained with a rabbit anti-mCherry (Takara-bio-clontech 632496, 1:500) and mouse anti-GFAP (Sigma G3893, 1:500) primary antibodies. After washing with 0.1 M PBS containing 0.1% Triton X-100, the sections were incubated with Alexa-488 and Alexa 594 conjugated secondary antibodies (Jackson Immunoresearch, 1:250). Fluorescent images of the mounted sections were obtained with a confocal microscope (LSM880, Carl Zeiss).

### Surgical preparation for electrophysiological recording in the trigeminal ganglion

Four to six weeks following AAV injections, animals were deeply anesthetized with urethane (1.5 g/kg, i.p.) and mounted on a stereotaxic frame. A homoeothermic control system kept the core temperature at 37.5–38°C. Animals were intubated and breathed O_2_-enriched room air spontaneously. Physiological parameters were continuously monitored (PhysioSuite, Kent Scientific, CapStar-100, CWE). A craniotomy was made to expose the cranial dura above the left visual cortex and the adjacent portion of the transverse sinus. Another small craniotomy was made over the right hemisphere to allow the insertion of the recording electrode into the left trigeminal ganglion. The exposed dura was bathed with a modified synthetic interstitial fluid ^24^.

### In vivo single-unit recording of trigeminal meningeal nociceptors

Single-unit activity of meningeal nociceptors (1/rat) was recorded from their trigeminal somata using a platinum-coated tungsten microelectrode (50–100 kΩ; FHC) ^24^. Meningeal nociceptors were identified by their constant response latency to electrical stimuli applied to the dura above the ipsilateral transverse sinus (0.5 ms pulse, 1-3 mA, 0.5 Hz). We recorded the activity of Aδ (1.5 ≤ CV ≤ 5 m/s) and C nociceptors (CV < 1.5 m/s). Nociceptor activity was digitized and sampled at 10 kHz using power 1401/Spike 2 interface (CED). A real-time waveform discriminator (Spike 2, CED) was used to create a template for the action potential, which was used to acquire and analyze nociceptor activity.

### Assessment of changes in meningeal nociceptor mechanosensitivity

Mechanical receptive fields of meningeal nociceptors were first identified using von Frey monofilaments ^24^. We then used a servo force-controlled mechanical stimulator (Series 300B, Aurora Scientific) for quantitative changes in mechanical responsiveness. Mechanical stimuli were delivered to the dura using a flat-ended plastic probe (0.5 or 0.8 mm) ^24^. Two ramp-and-hold stimuli were applied in each trial (rise time, 100 msec; stimulus width, 2 sec; interstimulus interval, 120 sec) and included a threshold stimulus (typically evoking at baseline 1–2 Hz responses) followed by a suprathreshold stimulus (2 to 3X of the threshold pressure). Stimulus trials were delivered every 15 min ^24^. Ongoing discharge was recorded between the stimulation trials. Basal responses to mechanical stimuli were determined during at least four consecutive trials before the chemogenetic stimulation.

### Recording changes in direct current potential and cerebral blood flow

To verify that chemogenetic manipulation of cortical astrocytes does not produce CSD ^14,15,20,21^, we recorded changes in DC potentials and cortical blood flow (CBF) in a subset of experiments. For DC recordings, a glass micropipette (70–120 kΩ) was implanted ∼1 mm below the pia overlying the visual cortex. DC potential was recorded (EXT-02F, NPI Electronic) using a reference Ag/AgCl ground electrode (A-M Systems) placed in the right temporal muscle. CBF changes were recorded using laser Doppler flowmetry (Vasamedic) ^24^. DC and CBF data were continuously sampled via an A/D interface (micro1401 and spike 2 software, CED, Cambridge, UK).

### Assessment of cephalic tactile pain hypersensitivity

Behavioral testing was performed as previously described ^26,45,46^. Animals were placed in a flat-bottomed acrylic apparatus (20.4 cm x 8.5 cm) and left to habituate for 15 minutes. To determine if animals developed pericranial tactile hypersensitivity, the skin region, including the midline area above the eyes and 2 cm posterior, was stimulated with von Frey (VF) filaments (0.6–10 g, 18011 Semmes-Weinstein kit). We evaluated changes in withdrawal thresholds and non-reflexive pain responses to stimulation by recording four behavioral responses, including: 0) *No response*: The rat did not display any response to stimulation 1) *Detection*: The rat turned its head towards the stimulating object and explored it, usually by sniffing; 2) *Withdrawal*: rat turned its head away or pulled it briskly away from the stimulating object (which usually followed by scratching or grooming of stimulated region); 3) *Escape/Attack*: rat turned its body briskly in the holding apparatus to escape the stimulation or attacked (biting and grabbing movements) the stimulating object. Starting with the lowest force, each filament was applied 3 times with an intra-application interval of 5 seconds, and the behavior that was observed at least twice was recorded. The force that elicited three consecutive withdrawal responses was considered the threshold, and the recorded score was based on the most aversive behavior noted. A cumulative response score was determined by combining the individual scores (0–3) for each one of the VF filaments tested.

### Open-field test

Open field behavior was monitored in a 43 × 43 × 30 cm arena illuminated with a white LED (80 lux) using Activity Monitor (Med Associates) ^47^. The system evaluates the movement of animals in the horizontal and vertical planes. We assessed the total distance moved and vertical activity (rearing) in each session for 20 minutes. A decrease in locomotion in the predesignated center zone was also assessed as a measurement of anxiety.

### Drug treatments

We activated the Gq-coupled hM3dq using an intraperitoneal injection of 1 mg/kg water-soluble clozapine N-oxide (CNO, Hello Bio) ^15,21^. CGRP signaling was blocked using a murine anti-CGRP mAb. A matched murine isotype control antibody was utilized as the control treatment. Antibodies were provided by TEVA Pharmaceutical Industries Ltd. and formulated in phosphate-buffered saline (PBS). Anti-CGRP mAb and the isotype control were injected i.p. at 30 mg/kg ^45,46,48^ 48 h before astrocyte stimulation.

### Statistical analysis

Data were analyzed and plotted using Prism 10 software and are presented as the median and interquartile range (IQR). Group data are plotted as scatter dot plots, with the median and interquartile range. Differences in nociceptor activation and sensitization propensities were analyzed using a two-tailed χ^2^ test. All other data were analyzed using the non-parametric two-tailed Mann-Whitney, Wilcoxon, or Friedman tests with corrections for multiple comparisons by controlling the false discovery rate. p, q < 0.05 were considered significant. Criteria used to assess meningeal nociceptor activation and sensitization were based on our previous studies ^24,25^. P-values are indicated as follows: p < 0.05 (*), p < 0.01 (**), and p < 0.001 (***).

## Acknowledgments

We thank members of the Levy lab for helpful discussions. The study was supported by NIH grants: R21NS101405; R01NS086830; R01NS078263; and R01NS115972 to DL. Support was also provided as part of an academic research collaboration between the Levy Lab at Beth Israel Deaconess Medical Center, Harvard Medical School, and Teva Pharmaceutical Industries Ltd.

## Author contributions

DB, JZ, JS, and DL contributed to the conception and design of the study; DB, JZ, and DL contributed to the acquisition and analysis of data; DB, JS, and DL contributed to drafting the text or preparing the figures.

---

**Supplement Figure S1.**
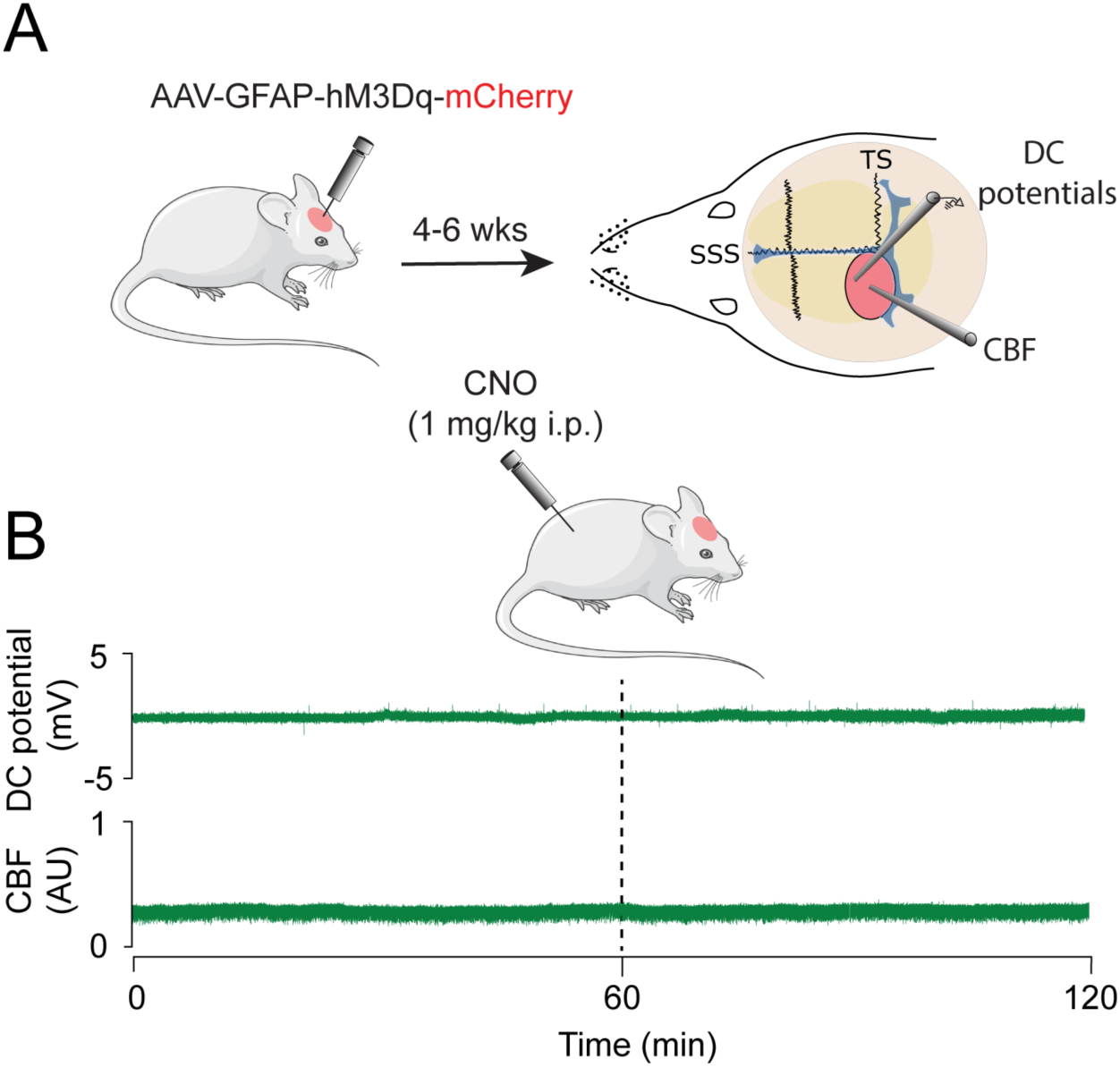
Chemogenetic activation of visual cortex astrocytes Gq-GPCR signaling does not trigger cortical spreading depolarization. **A.** Experimental design for recording direct current (DC) potentials and cerebral blood flow (CBF) to assess CSD triggering following chemogenetic cortical astrocyte Gq-GPCR stimulation. DC potentials and CBF were recorded via a skull opening above the visual cortex injected with AAV5-GFAP::hM3Dq-mCherry. **B**. An example of a two-hour experimental trial showing no change in DC potential or alteration in CBF indicative of CSD in the 1 h following injection of CNO (1 mg/kg, i.p.) in hM3Dq^(+)^ rats.

